# Contextual Influences on Value-based Choice

**DOI:** 10.1101/340695

**Authors:** Vincent Man, Chloe Kovacheff, William A. Cunningham

**Affiliations:** Department of Psychology University of Toronto 100 St. George Street Toronto, Ontario M5S 3G3 Canada; Rotman School of Management University of Toronto 105 St. George Street Toronto, Ontario M5S 3E6 Canada

## Abstract

Biases in choice behavior are shaped by both immediately encountered cues as well as the background context in which these cues are embedded. Here we examine the mechanisms that underlie the integration of contextual and cue information, and the manner in which these sources of information shape behavior. We demonstrate contextual influence on choice dissociated from cue information using a value-based multi-alternative choice task while recording neural activity using electroencephalography. Consistent with work on negativity biases, we show that contextual attributes related to loss, reflected along central-parietal sites in the gamma (30-59 Hz) frequency range, are prioritized and drive behavior to prevent loss. We employ a multi-alternative sequential sampling computational model to show that contextual and cue information are integrated through the decision process to shape choice responses, and link this integrative process to a neural signature in the gamma frequency band.

## Introduction

Adaptive behavior in the world necessitates integrating relevant information from multiple possible sources and exploiting this information to inform decision processes. Two potential sources of relevant information include the immediate cues encountered in the environment, and the properties and contingencies of the environment itself. A foraging animal that encounters a relevant cue, such as a bush that might hold ripe fruit, is more likely to survive if it integrates decisions to approach with knowledge about the season and presence of potential predators. Integrating this contextual knowledge becomes critical for correctly evaluating the trade-off between predatory threat and opportunity for food. Here we ask how these sources of information might be differentially processed in the brain, as well as the nature of the computations that allow background contextual and immediate cue information to be integrated for decisionmaking.

While recent research has placed emphasis on the hierarchical organization between context and cues, our current focus aims to address the open question of how different stages across the hierarchy shape one another; that is, on the *integrative* processes between contextual and cue information. Such a hierarchical organization, in which immediate cues are embedded within higher-order structure with shared statistical features (Botvinick, 2012; Collins and Frank, 2013; Gershman et al., 2010), is a useful framework to adopt as it decomposes the study of contextual influence on decisionmaking into tractable problems of contextual representation versus control. Problems of representation concern the consolidation of shared statistical features across a given environment (Tse et al., 2007; Wilson et al., 2014), and is largely influenced by the literature on cognitive maps (Tolman, 1948; Gershman et al., 2010). On the other hand, by control we emphasize the processes that use contextual information for the purpose of guiding responses to immediately encountered information (Cohen et al., 2000; Daw et al., 2011). The latter problem is the focus of the current inquiry.

In many ways this hierarchical relationship between context and cue mirrors the organization of neural systems across the brain (Badre and D’Esposito, 2009; Ribas-Fernandes et al., 2011). Specifically, there has been increasing evidence that higher-order contextual structure might be represented across the prefrontal (Schuck et al., 2016) and medial temporal cortices (Burgess et al., 2002; Diana et al., 2007). There has also been much work on how the brain supports control over behavior, with emphasis on the role of the anterior cingulate cortex (Behrens et al., 2007; MacDonald et al., 2000), as well as distributed regions across frontoparietal cortices (Nieuwenhuis et al., 2004), in particular relation to attentional processes (Niv et al., 2015). Electrophysiological experiments have demonstrated the importance of slow theta (Θ, 1-7 Hz) oscillations along the frontal midline for both memory maintenance (Hsieh and Ranganath, 2014) as well as control (Cavanagh and Frank, 2014; Summerfield and Mangels, 2005). Additionally, fast gamma (γ, > 30 Hz) oscillations across the cortex, and particularly in a frontoparietal system, has been linked to computations driving value-based choice (Polanía et al., 2014), and support the binding between contextual representations and incoming cue-(item) level information (Jensen et al., 2007; Nyhus and Curran, 2009). These findings provide important guidance for delineating the component processes that drive context-dependent choices; however, specifying the potential mechanistic pathways through which these neural systems give rise to behavior remains an important challenge.

We address this open challenge by examining how the brain supports higher-order contextual information that is dissociated from cues, as well as how precise neural signatures and computational mechanisms account for the integration of contextual- and cue-level information. We probe both the isolated and interactive processes across contexts and cues by testing participants on an adaptation of a multi-alternative value-based forced-choice paradigm (Mowrer et al., 2011) while recording neural activity using noninvasive electroencephalography (EEG). We exploited the high temporal resolution and spatial breadth of this neuroimaging modality to probe whether different contextual attributes might be represented in parallel neural traces across the frequency spectrum in a distributed fashion. Under our experimental paradigm (Figure 1), at a given moment the potential to acquire a gain or prevent a loss outcome is given by the incoming cue information. Importantly, we include a response time limit to induce speed-accuracy trade-off processes; this allows us to examine the response profile of error responses and consequently, whether strategic attention shifts towards contextually advantageous choice options. We manipulate the magnitude of the potential monetary outcome across task blocks to shape the context in which these trial-level cues are encountered. This dissociation between the information conferred by the context and by the cue allows us to employ an analytic strategy that disentangles sustained, or tonic contextual influence from context-cue interactions. We do so behaviorally by examining response bias towards contextually-favored choice alternatives on trials in which the cue indicates no monetary consequence. We take a similar approach in dissociating context-unique neural markers by examining oscillatory signal that vary across contextual states, and isolating contextual components of the resulting signal.

**Figure 1.**
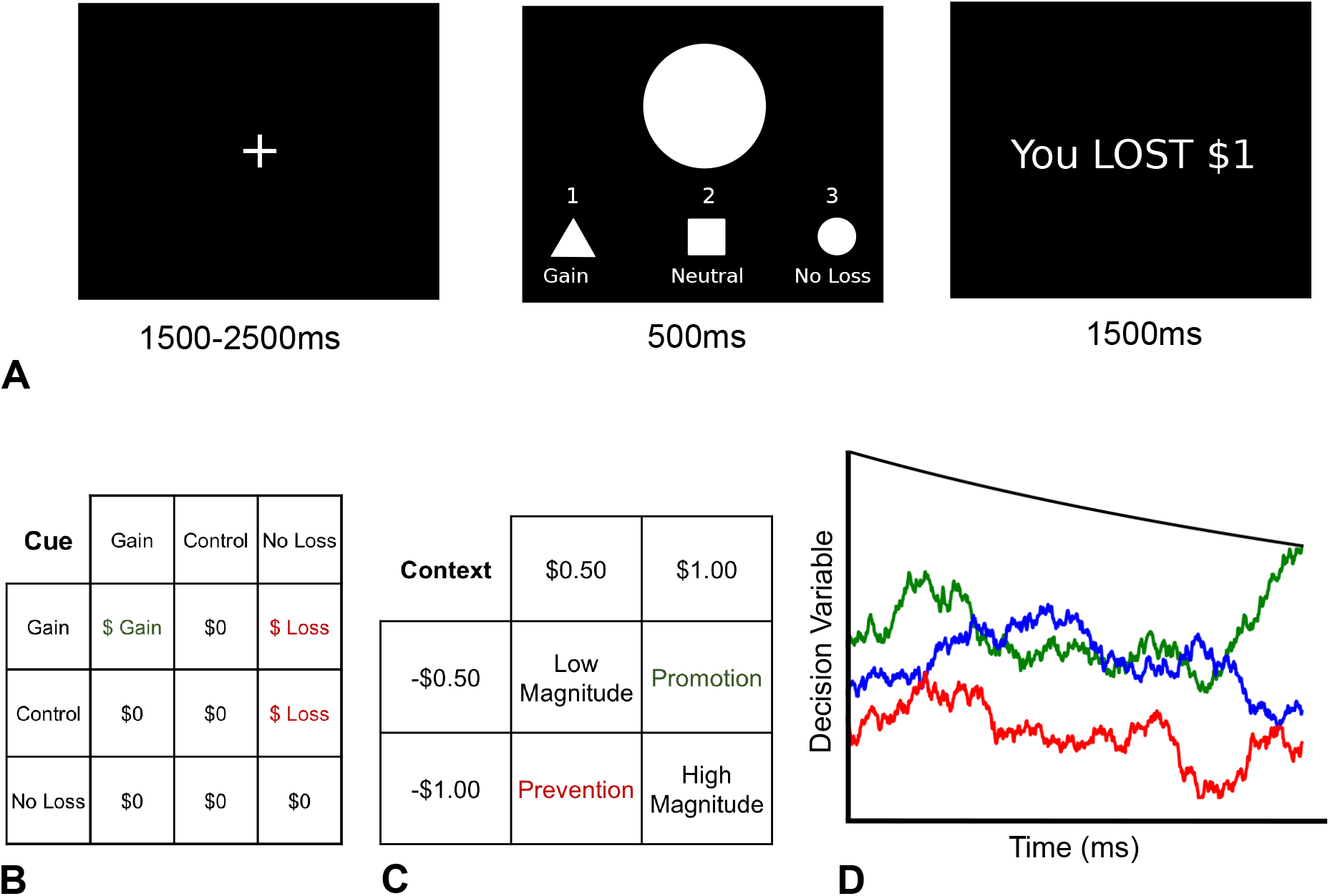
Experimental paradigm and computational model. A) Sequence of events within a trial (see supplementary methods for details) B) Matrix of potential outcomes given the presented cue (columns) and potential responses (rows). C) Context was manipulated via outcome, with contextual gain values of +$0.50 and $1.00 (columns) and contextual loss values of −$0.50 and −$1.00 (rows). D) Schema of the multi-alternative sequential sampling model, with three noisy accumulators for each choice alternative. The first accumulator to reach the collapsing upper boundary gives the behavioral choice for that trial, with the reaction time given by the corresponding value on the Time (x-) axis.

To address the integrative processes between contextual and cue-level information and how these processes drive choice behavior, we tested a potential computational mechanism and probed the nested trial-level neural signal along spatial, temporal, and oscillatory frequency dimensions. We tested the possibility that the context- and cue-level information is integrated moment-to-moment in the decision process, and that this integrated signal explains observed contextual influence on choice behavior. We also tested an alternative mechanism by which observed context-dependent choices need to additionally recruit an initial representation of the context, which sets the stage for later integrative processes (see Bornstein et al., 2017; Lositsky et al., 2015; Mulder et al., 2012). To test these potential mechanisms of context-cue interactions computationally, we built upon an approach that frames decision processes as an evidence accumulation problem using sequential sampling models (SSMs; (Brown and Heathcote, 2008; Bogacz et al., 2006; Ratcliff, 1978; Roe et al., 2001; Usher and McClelland, 2001; Wald, 1947). These models are unified by the principle that observed behavior in decision problems is driven by the noisy accumulation of evidence respective to each alternative. SSMs importantly capture speed-accuracy trade-offs in behavior, and have been successfully applied to explain decisions across perceptual (Gold and Shadlen, 2001; Tavares et al., 2017) and value-based (Hunt et al., 2012; Hutcherson et al., 2015; Krajbich and Rangel, 2011; Mormann et al., 2010; Polanía et al., 2014) domains. Importantly, both their development and application have been closely linked to underlying biology, across neural modalities (Hare et al., 2011; Hunt et al., 2012; Polanía et al., 2014; Shadlen and Newsome, 2001).

This link to underlying biology is particularly important given that both computational predictions describe context-cue integration as a process that unfurls over the decision period. As such, neural traces reflecting this integrative process should occur after both contextual and cue-level information is available and within the span of a decision trial, and relate to the relative activation of the evidence-accumulating model mechanisms. To test this prediction, we specified contrasts that specifically probed statistical interactions between contextual features and cue information, using an out-of-sample approach to first delineate the spatiotemporal topology of any signature supporting any interaction, then time-frequency properties of this signature at its relevant brain sites.

Extending our inquiry into how context and cue bind together, we further capitalized on our paradigm and analytic strategy to examine whether certain contextual attributes (i.e. loss versus gain features of the context) might be prioritized over others. We adopt the perspective that a model of the external context is internalized in the form of a goal, in that they are placed in relation to concrete objectives and associated with potential action strategies (Dolan and Dayan, 2013; Rushworth et al., 2011; Solway and Botvinick, 2012). Importantly here, goals are shaped by external contingencies: our foraging animal might have attained through experience the knowledge that the current season invites predators to an area. As a result, goals to prevent harm outweigh goals to exploit foraging opportunities, and it executes a congruent action set by seeking shelter instead of approaching the bush. This perspective allows us to ask whether certain types of goals are prioritized by default over others. For example, given equal likelihood between the presence of abundant food or predators at a certain patch, our foraging animal may default towards prevention goals and again seek shelter because the threat of harm intrinsically outweighs potential opportunity costs (Higgins, 1997). To the extent that certain environmental demands take precedence with regards to attentional and motivational resources for survival, the goals respective to those environmental states should be more heavily weighted.

## Results

### Preferential response to value

We conducted an initial investigation with an independent sample that completed a simple non-contextual version of the three alternative forced-choice task (described in supplementary methods). The match-to-sample nature of the task means that participants are able to succeed at the task by only attending to the presented cue and choosing the matching response, without representing any associated reward outcomes. If participants solely focus on matching response to the presented cue, we expect the associated value of the *gain* and *no loss* choice alternatives to be irrelevant and thus similar response tendencies across all three choices.

Instead, we found that participants showed preferential response to the value-based *(gain* and *no loss)* cues. We found significantly higher accuracy for gain (b=1.475, SE=0.112, z=13.196, p<0.001) and no loss (b=1.211, SE=0.106, z=11.385, p<0.001) cues, compared to control (see Figure 4a). In this same model we controlled for potential interactions with the between-subjects outcome magnitude assignment (see supplementary methods), and found no significant main effect of outcome magnitude assignment (χ^2^(1)=0.444, p=0.505) nor significant interaction between the cue type and outcome magnitude (χ^2^(2)=3.130, p=0.209).

Reaction time (RT) analyses also confirmed that participants attended to valued cues preferentially over the control alternative, despite explicit instructions to match shapes on all trials. We found a significant main effect of the cue condition (F(2,5394)=74.363, p<0.001; see Figure 4b), after controlling for the main effect of the between-subjects outcome magnitude assignment and for interactions between cue and outcome assignment. There was no significant main effect of the outcome magnitude assignment (F(1,55)=0.460, p=0.501) or its interaction with the cue type (F(2,5394)=0.836, p=0.434).

### Context-dependent behavior

In an independent sample we recorded 64-channel EEG data from healthy adults performing a contextual version of our multi-alternative three-alternative forced choice task (see supplementary methods). We replicated the effect that participants respond preferentially to valued cues, with significantly higher accuracy for gain (b=1.455, SE=0.040, z=36.55, p<0.001) and no loss (b=1.413, SE=0.040, z=35.62, p<0.001) cues, compared to control. Confirming our hypothesis that contextual demands can shift the focus towards appropriate cue types, a model that includes the contextual condition as a higher-level moderator of cue condition had a better fit to data (χ^2^(9)=26.571, p=0.002), compared to a nested variant that didn’t include the context regressor. We found a significant effect of the *no loss* cue in the *prevention* context (in which focus on preventing loss is optimal) in predicting response accuracy (b=0.414, SE=0.112, z=3.702, p>0.001; Figure 2a).

**Figure 2.**
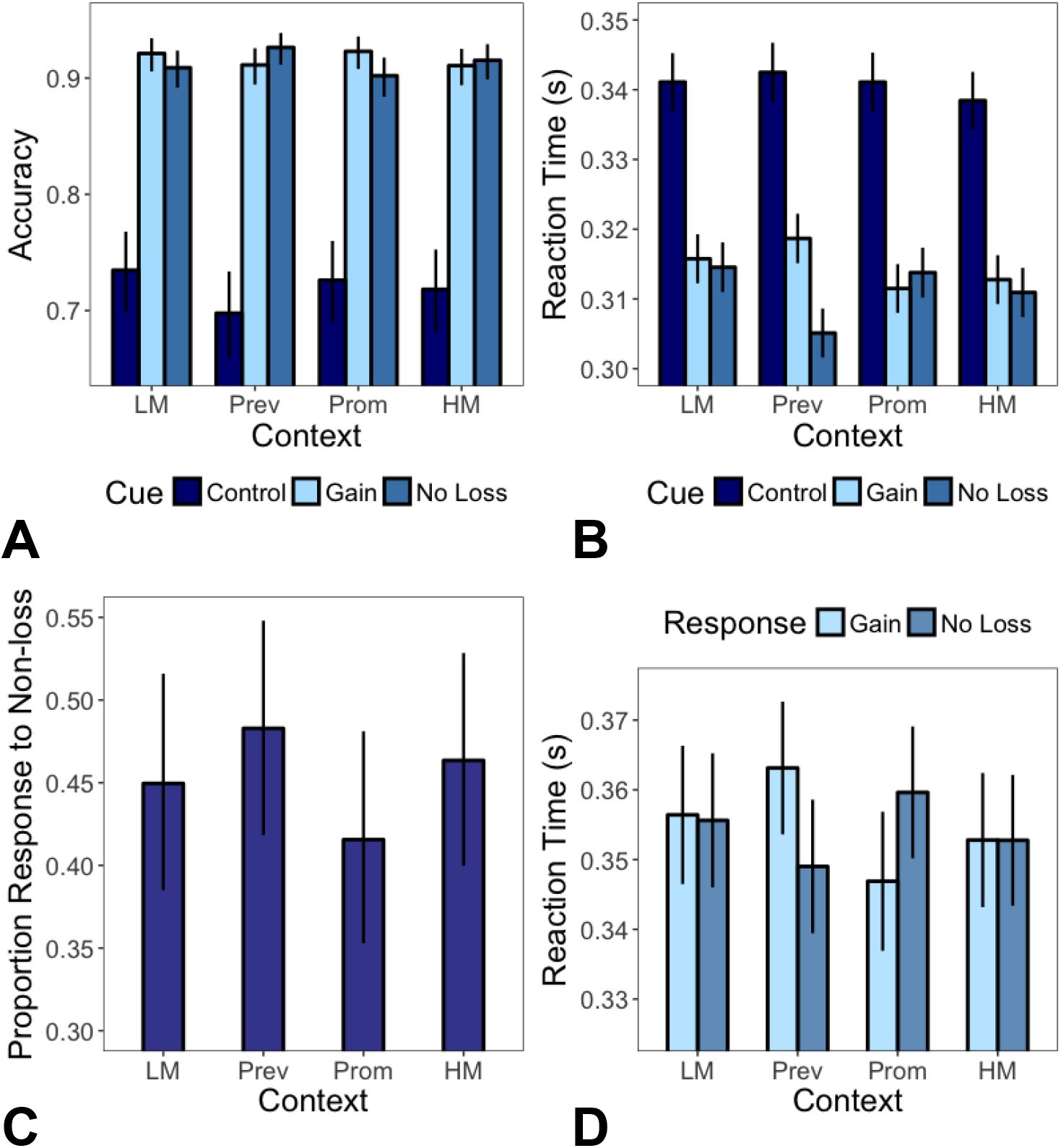
Behavioral contextual influence. Participants show greater response accuracy (A) and faster reaction times (B) for valued, compared to non-valued, cues. Participants were significantly more accurate and faster for no loss cues in the prevention context. Error responses on trials with the non-value control cue demonstrate effects of contextual influence: participants showed more responses (C) and faster reaction (D) to the no loss cue in the prevention context, and for the gain cue in the promotion context. The proportion response (C) effect was not significant but the reaction time (D) effect was significant. Bars represent estimated means ± 95% CI. LM: Low magnitude; Prev: Prevention; Prom: Promotion; HM: High Magnitude.

RT analyses again provided convergent evidence that participants attended to valued choices preferentially over the control alternative, despite explicit instructions to match shapes on all trials. We found a significant interaction between cue and context (F(6,20151)=12.76, p<0.001). Simple effects tests showed that prioritized response was moderated by context such that participants responded more quickly to the no loss cue in the prevention context (b=0.007, SE=0.001, t(19670)=9.448, p<0.001). We also found an effect of gain cue prioritization in the promotion context (b=-0.001, SE=0.001, t(19710)=-1.986, p=0.047) (Figure 2b).

To the extent that individuals represent contextual properties beyond cue-based reward outcomes, we expected error response biases to be strategically aligned with current contextual demands, and exhibited throughout the block. Given that context shapes the outcome magnitudes for the valued cue trials, if contextual demands impose tonic influence on behavior, error responses on the non-valued control cue trials should align with the advantageous strategy in the current context.

We tested this third critical behavioral hypothesis by modelling behavior as a function of the type of error response (i.e. to gain or no loss) and context, only on trials with control, non-valued, cues. We found that participants exhibited bias in their error responses in a context-congruent manner, consistent with our hypothesis (Figure 2c), though the effect was not significant (χ^2^(3)=5.001, p = 0.172). However, we found a significant interaction effect (F(3,2531)=4.561, p=0.003) of RT: simple effects tests demonstrated significantly faster RT for erroneous non-loss responses to control cues in the prevention context (b=0.014, SE=0.005, t(2526)=2.743, p=0.006) and significantly faster RT for erroneous gain responses to control cues in the promotion context (b=-0.013, SE=0.005, t(2630)=-2.424, p=0.015) (Figure 2d). Together this pattern of context-dependent error responses reveals contextual influences dissociated from cue-driven behavior.

### Neural signatures of context

If there are distinct neural mechanisms that underlie higher-order processes, probing the FFT signal with a set of orthogonal contrasts that controls for cue level variability should result in distinct patterns that correspond to contextual attributes without an effect of cue. Otherwise, if the choice biases are driven only by cue-related processes, we would expect no effect across all contrasts. We specified a linear mixed-effects model (described in the supplementary methods) that allowed us to separately look at how the brain processes contextual gain, contextual loss, and the informativeness of a context. Critically, we exploited our two control context conditions, along with the balanced experimental design that necessitated the same number of cue types among trials within a context, to probe a specific contextual feature (e.g. contextual loss) while controlling for variability at the cue level (e.g. the amount of potential gain given by the trials).

An area covering midline central-parietal sites in the low-γ (30-59 Hz) band showed significant effects for contextual loss features (F(1,261)=5.612, p=0.019; Figure 3a), but not for contextual gain (F(1,261)=0.020, p=0.887) or informativeness (F(1,261)=0.069, p=0.793). Frontal electrodes in the Θ (4-7 Hz) band also showed significant effects for contextual loss features (F(1,261) = 10.077, p = 0.002; Figure 3b) but no effects of gain (F(1,261)=0.010, p=0.757) or informativeness (F(1,261)=0.049, p=0.825). Finally, parietal electrodes in the Δ (1-3Hz) band also showed significant effects for contextual loss (F(1,261)=9.864, p=0.002; Figure 3c) but no effects of gain (F(1,261)=0.035, p=0.851) or informativeness (F(1,261)=0.003, p=0.957). While we found three neural signature across frequency space reflecting greater contextual loss processing, we did not find any effect for the contextual gain or informativeness contrasts (see Figure S2a).

**Figure 3.**
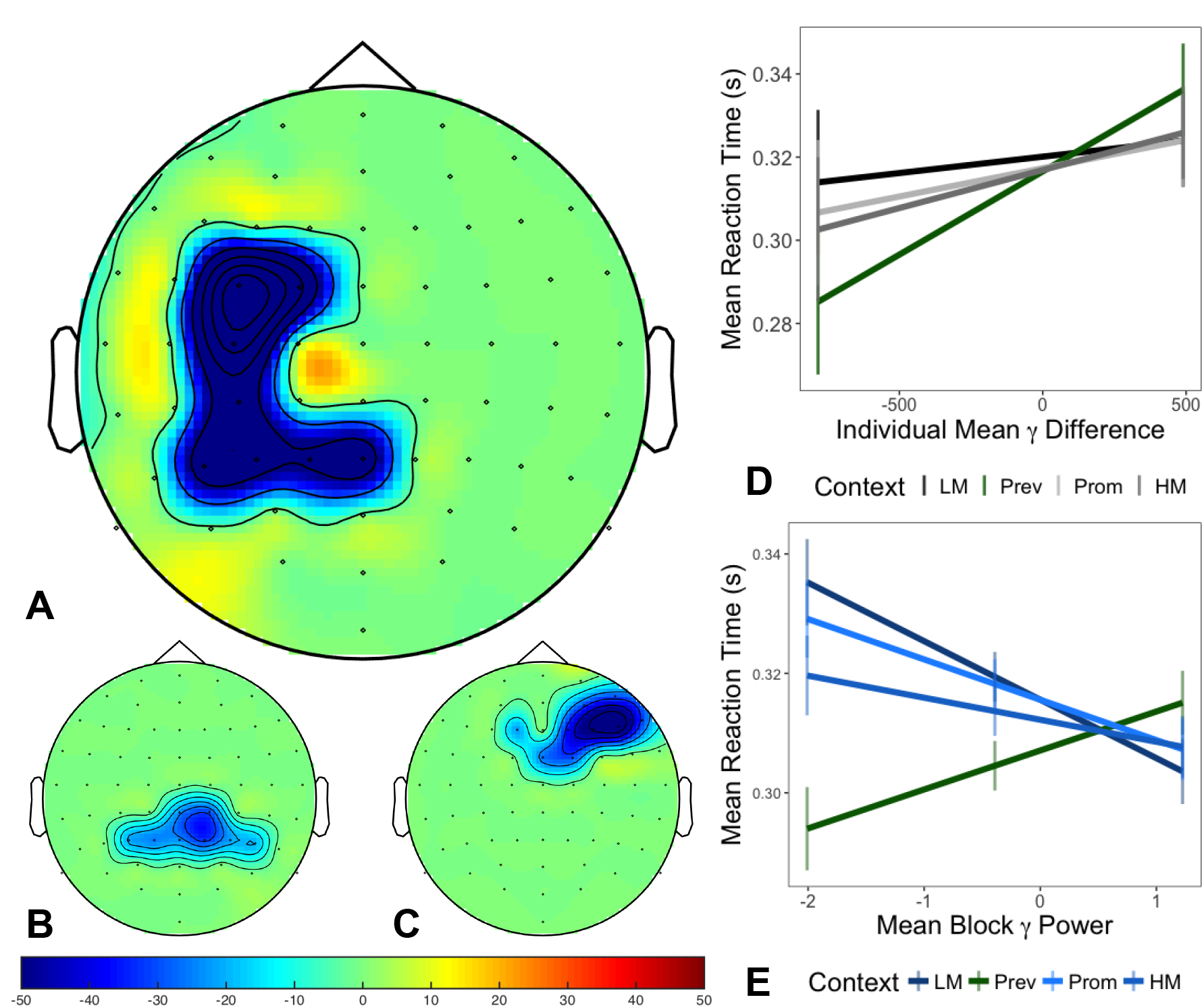
Contextual loss representation. A) Significant channels for the low-γ band. B) Significant channels for the Δ band. C) Significant channels for the Θ band. D) Individual differences in low-γ power predicted faster responses to the no loss cue under the prevention, but no other, context. E) Within-participant low-γ power predicted faster reaction times to the prevention context, but no other, context. Lines represent estimated means ± 95% CI.

If this neural emphasis on loss features reflects contextual processing, we expected that it should correspond to observed behavioral biases related to loss. Our reasoning was that neural signatures of contextual processes should have downstream influence on behavior. Since it is more advantageous to more strongly represent loss-related contextual attributes under the *prevention* blocks, the frequency signatures described above should predict behavior to prevent losses more so within these *prevention* blocks.

We tested this in two ways. First, we looked at whether individual differences in this neural signature across contextual conditions drove performance on the task (see supplementary methods). We found a significant two-way interaction between context and individual differences in mean low-*γ* power differences (according to the Loss Features contrast across central-parietal channels; Figure 3a) in predicting mean RT (F(3,1012)=17.537, p<0.001, Figure 3d). Specifically, we found that individuals with decreased low-*γ* power demonstrated generally faster reaction times to successful choices, particularly in the *prevention* context. This was supported by significant simple main effects of mean *γ* difference across participants for the *prevention* context (b=0.011, SE=0.005, t(98)=2.267, p=0.026) compared to the other contexts *(promotion:* b=0.006, SE=0.005, t(97)=1.184, p=0.239; *high magnitude:* b=0.006, SE=0.005, t(97)=1.301, p=0.196; *low magnitude:* b=0.001, SE=0.006, t(97)=0.229, p=0.819).

Furthermore, if this central-parietal *γ* power constitutes a neural representation at the high-order contextual level, this signal should have a tonic influence on behavior to prevent loss throughout the span of the *prevention* block (i.e., even during trial in which the *no loss* cue is not presented). We tested whether RT is predicted by the interaction between this neural signature and task condition. Critically for this analysis, the block-level neural regressor was computed by only averaging across *gain* and *control* cue trial values of mean *γ* power across the central-parietal sites in Figure 3a, but was used to predict mean RT only across *no loss* cue trials (see supplementary methods). In other words, this neural regressor constitutes variation in representing contextual loss features, but was computed without inclusion of cue-level loss trials, thereby dissociating context and cue while controlling for neural responses to stimulus presentation. Within participants, we found that block-level *γ* power across central-parietal sites significantly interacted with the contextual condition to predict mean RT to no loss cues (F(3,7998) = 22.393, p < 0.001; Figure 3e), even after controlling for neural signal in pre-trial fixation periods. Specifically, we found that within participants, decreased mean *γ* power predicted faster responses to *no loss* cues only within the *Prevention* context (b=0.006, SE=0.007, t(80)=8.597, p<0.001, R2 = 0.009, Cohen’s d = 0.192), whereas for the other contexts, the decreased mean power did not drive faster reaction times to *no loss* cue trials.

### Computational mechanisms driving contextual influence

We probed two potential computational pathways by which context exerts influence on decision processes. By one hypothesis, the integration of contextual information into the moment-by-moment accumulation process can capture the observed behavioral context effects in cue match accuracy and RT. Alternatively, the current context might additionally initialize the decision process; this implies that the identity of the current context needs to be represented prior to decision accumulation process (Bornstein et al., 2017; Lositsky et al., 2015; Mulder et al., 2012).

We formalized these hypotheses with a multi-alternative variant of a SSM (m-SSM; Figure 1d; see supplementary methods for the formal model and details), similar to previous approaches (Churchland et al., 2008; Krajbich and Rangel, 2011; Tsetsos et al., 2011), and according to specific features of our experimental design. Critically, the stochastic accumulator is driven by both the cue information formalized with the cue boost parameter *c*, as well as the (normalized) outcome magnitudes of all available alternatives, which in our experiment is the contextual information. This design importantly allowed us to disentangle the contribution of cue processes (e.g., the response matching component of our paradigm) with the processing of value across all choice alternatives. We tested a divisive normalisation mechanism (Louie et al., 2011; 2013) to account for choice set effects, or the local context of an alternative against other available options (Tversky and Simonson, 1993). Importantly, divisive normalisation includes a weight parameter *w* that determines the degree to which objective outcome magnitudes are shaped by the choice set.

We used the independent sample of the initial non-contextual experiment to validate this design for the m-SSM by testing against other model variants, using the group data out-of-sample. The models differed along two components: we modified either the core value transformation driving the accumulator, the starting position of each accumulator, or both (Table S1). Our criterion was that the model predictions should fall within the 95% CI of the empirical data across all conditions. Specifically, a good fitting model should reproduce the relative differences in accuracy and RT between the valued *(gain* and *no loss)* and non-valued *(control)* cues, and approximate the means of each condition in the empirical data.

Two model variants, including the m-SSM, were the only models whose out-of-sample predictions fell within the 95% CI of the empirical data across cue conditions, replicating the accuracy (Figure 4c; Models 1 [red line] and 3 [yellow]) and RT (Figure 4d) effects, for both the *Low Magnitude* (LM) and *High Magnitude* (HM) between-subject assignments. We favored the m-SSM because modelling the starting values linearly (Table S1) allows us to scale the model more efficiently, using less parameters (see supplementary methods). The divisive normalisation of choice alternative values critically allowed the m-SSM to reproduce the response accuracy patterns observed in the behavioral data: we found significant higher accuracy for gain (b=1.239, SE=0.142, z=8.700, p<0.001) and no loss (b=1.157, SE=0.143, z=8.101, p<0.001) cues, compared to control (Figure 4a), after controlling for potential interactions with the between-subjects outcome magnitude assignment (Main effect of outcome magnitude: χ2(1)=0.043, p=0.835; Interactions between outcome magnitude and cue: χ2(2)=0.080, p=0.961). Alternative model variants with outcome magnitudes transformed subtractively did not capture these accuracy effects (Figure 4c-d and Table S1: Models 4 [green line], 5 [blue line], and 6 [purple line]).

**Figure 4.**
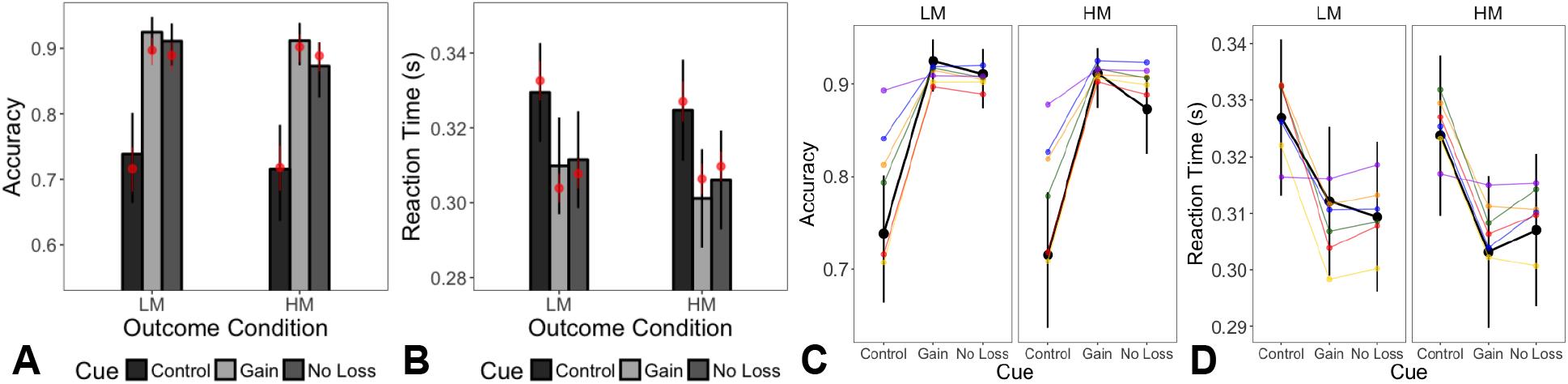
Behavioural psychometrics and model comparison for Experiment 1. The grayscale bars show logistic (A) and linear (B) mixed-effects model estimates from the empirical data on left-out even trials, with lines depicting 95% CI. The red bars show the m-SSM out-of-sample predictions. C and D) The empirical data (even trials) are depicted with the point ranges in black, with points representing estimated means and lines representing 95% CI from the logistic (C) or linear (D) mixed effects models. m-SSM predictions (fit on odd trials) are depicted as color lines: red (Model 1), orange (Model 2), yellow (Model 3), green (Model 4), blue (Model 5), purple (Model 6). Model 1 and 3 were the only models whose out-of-sample predictions fell within the 95% CI of the empirical data across cue conditions.

We also specified unique starting position for each accumulator within the m-SSM. This model feature was critical for reproducing the reaction time effects observed in the behavioral data: we found a significant main effect of the cue condition (F(2,3039)=57.993, p<0.001) with faster responses to the gain and no loss cues (Figure 4b), after controlling for potential interactions with the between-subjects outcome magnitude assignment (Main effect of outcome magnitude: F(1,3039)=0.093, p=0.760; Interactions between outcome magnitude and cue: F(2,3039)=1.623, p=0.197). Notably, an alternative model with a shared starting position across accumulators was not able to capture RT difference between the valued and non-valued cues (Figure 4d: Model 6 [purple line]).

We formalized the alternative hypothesis which posits that the identity of the current context additional initializes the decision process with a contextual model variant (cm-SSM; see supplementary methods for the formal model and details), which builds on the m-SSM by allowing unique accumulator starting positions between the *Promotion* and *Prevention* contexts. Both the m-SSM and the cm-SSM were able to capture key aspects of the observed behavioral data, out-of-sample (Figure 5). We favored the m-SSM given reduced model complexity; indeed, the best fitting parameters values for the two additional parameters of the cm-SSM (β_Prev_ and β_Prom_) were both 0.

**Figure 5.**
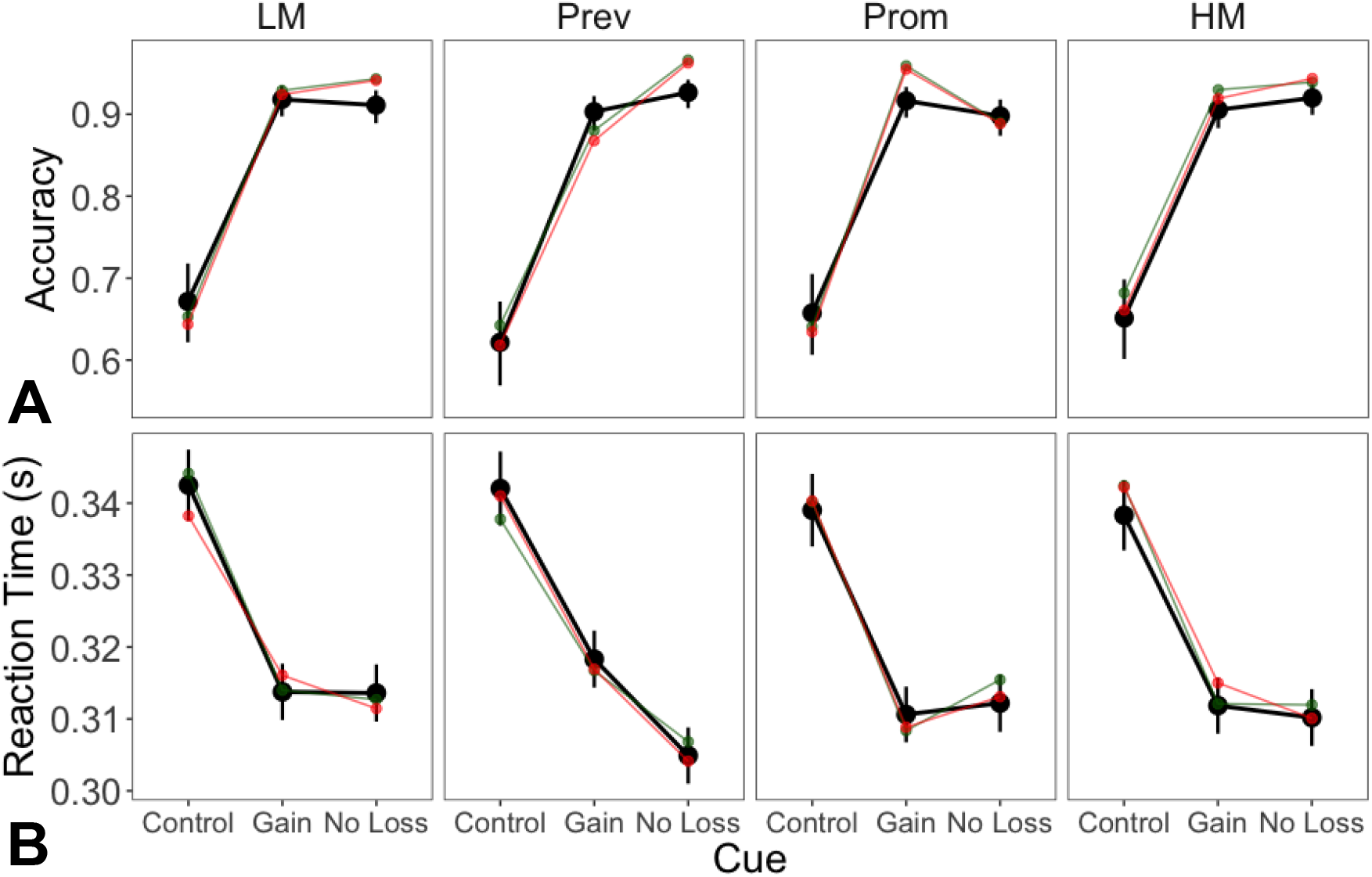
Model comparison for Experiment 2. The empirical data (even trials) are depicted with the point ranges in black, with points representing estimated means and lines representing 95% CI from the logistic (A) and linear (B) mixed effects models. Model predictions (fit on odd trials) are depicted with color lines: m-SSM (red) and cm-SSM (green). Note that the m-SSM has two fewer parameters than the cm-SSM (Table S2).

Critically, the m-SSM was able to reproduce context-dependent cue match accuracy rates for the key *prevention* and *promotion* contexts. Reproducing the empirical data, the contextual condition was a significant moderator of different accuracy rates across cue conditions (χ^2^(96)= 122.680, p<0.001; Figure 5a), with significant greater accuracy for the *no loss* cue (b=0.581, SE=0.211, z=2.749, p=0.006) and lower accuracy for the *gain* cue (b=-0.510, SE=0.168, z=-3.035, p=0.002) in the *prevention* context, and vice versa in the *promotion* context *(no loss* cue: b=-0.661, SE=0.182, z=-3.636, p=0.0003; *gain* cue: b=0.589, SE=0.195, z=3.013, p=0.003). The m-SSM was also able to produce the RT patterns in the empirical data: we found a significant interaction between cue and context (F(6, 11925)=8.171, p<0.001; Figure 5b), and simple effect tests demonstrated faster responses to the *no loss* cue in the *prevention* context (b=6.869, SE=0.939, t(11933)=7.315, p<0.001), and to the *gain* cue in the *promotion* context (b=-2.545, SE=0.933, t(11933)= −2.727, p<0.006).

### Neural signatures of context-cue interactions

Given our finding that context-cue integration occurs through the decision process, neural signatures reflecting integration should occur after both contextual and cue-level information are available. Further, we expected that such signatures should occur within the *γ* frequency range and along frontal or parietal sites, given that contextual traces predicting behavior occur in this range and space, described above, as well as previous work showing the role of frontoparietal *γ* in both context-item binding (Jensen et al., 2007; Nyhus and Curran, 2009) as well as evidence accumulation (Polanía et al., 2014).

We tested this by specifying a set of orthogonal contrasts that model the interaction between context and cue: specifically probing the contrast that compares context-congruent cue (e.g. *gain* cues in the *promotion* context) against incongruent cue trials (see supplementary methods). Using an out-of-sample approach, we found a set of central electrodes were significantly related to context congruency (Figure 6a). Further probing this congruency interaction contrast using the left-out trials, we found that this central site effect is shaped by increased low-*γ* power to context-congruent cues within a mid-trial period (Figure 6b).

**Figure 6.**
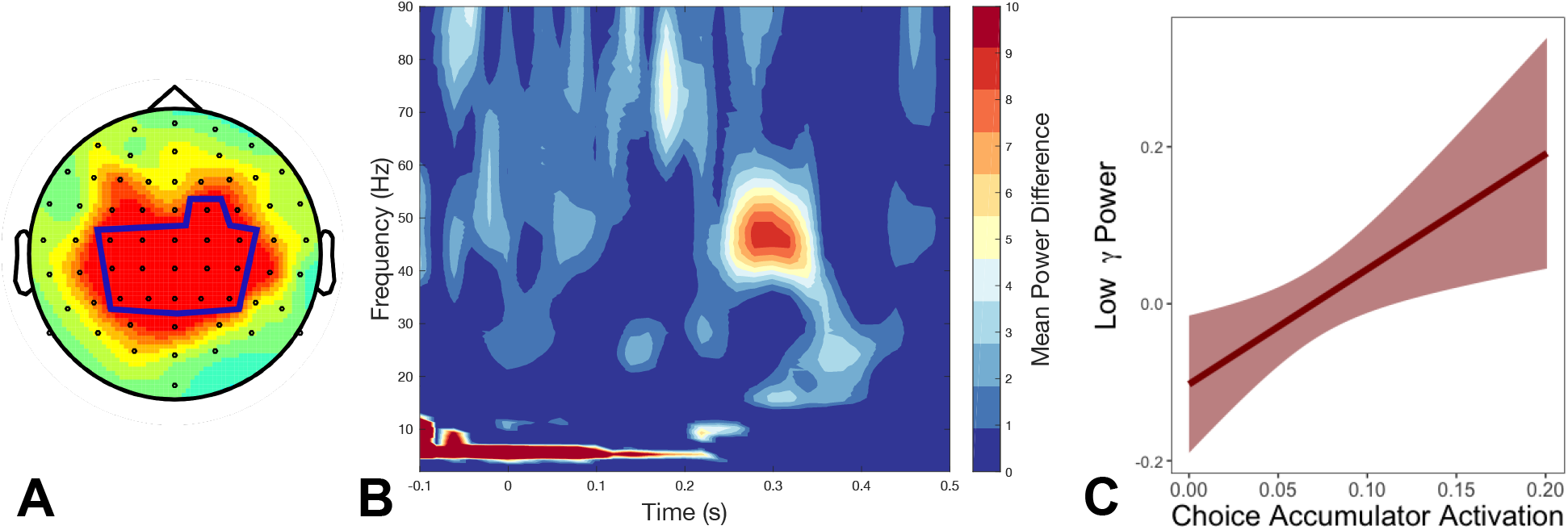
Mechanisms of context-cue interactions. Grayscale bars show logistic (A) and linear (B) mixed-effects model estimates from the empirical data. The red bars show the m-SSM out-of-sample predictions. The model was able to capture key experimental patterns in response accuracy (A) and RT (B). C) Topographical signature of context-cue interactions (showing 0.25 to 0.30 s post-stimulus; see Figure S3 for the full trial time span). D) Time-frequency analysis for the left-out trials, averaged across channels shown in the blue box in (C). Color heat demonstrates differences between the context-congruent and incongruent mean power differences. E) The extracted neural signal from the low-*γ* cluster in (C; 46-47 Hz, 280-300 ms) is significantly predicted by accumulator activation respective to the chosen alternative. Bars represent estimated means ± 95% CI.

Finally, to confirm that this neural signature is involved in integrative processes described by the computational model, we used individual participant model fits to compute the trial-by-trial activation of the model accumulator corresponding to the chosen alternative (see supplementary methods). We regressed the trial-by-trial neural signal, averaged over the constrained electrodes, time, and frequency windows (Figure 6b; using the peak difference at 48-49 Hz and 280-300 ms), onto the model-predicted accumulator activation while controlling for any effect of RT in a linear-mixed effects model. We found a significant positive relationship between this low-*γ* neural trace and the model accumulator activation (b=1.466, SE=0.545, t(33195)=2.689, p=0.007; Figure 6c).

## Discussion

To understand what influences adaptive behavior and decisions, it is necessary to examine the role of both proximal cues and the broader nature of the context itself. Here we addressed the open questions of how the brain supports contextual processes dissociated from the cues embedded within these contexts, and of how neural and computational mechanisms drive the integration of contextual information with cue-level information to shape choice behavior. Importantly, we exploited neural and computational analytic techniques, along with a value-based choice paradigm designed specifically to disentangle these component processes, to probe underlying choice strategies that are dependent upon contextual information.

As predicted, our results first demonstrated that context influenced how participants responded to cues. Critically, we found that speed-accuracy trade-off processes aligned with advantageous strategies in a given context: if behavior is solely driven by the identity of the cued alternative and its associated outcome magnitude (i.e. by only cue-level information), then we would expect no difference in behavior between errors toward the gain and no loss responses when the *control* cue is prompted. Instead, we found that under shifting contextual demands, participants demonstrated error biases and responded more quickly to context-congruent cues, even when the cue-level information was dissociated from value. This finding provides evidence of a separate and distinguishable influence of context over and above cue information.

Building upon these findings, we examined the neural correlates of contextual representations by employing an analytic approach that allowed us to dissociate the gain-, loss-, and information-related features of a context. This allowed us to address the larger question of how certain contextual attributes might be prioritized by default over others. If a certain contextual feature is weighted more heavily than another, we expected that neural systems supporting the representation of this contextual attribute might show stronger effects. We found that the contextual loss feature was more heavily represented across multiple spatial regions and frequency bands of neural oscillation. We found this contextual loss effect in frontal Θ, parietal Δ, and central-parietal low-*γ* traces.

Importantly, we represented the neural signature of context completely independently of cues by regressing behavior associated with *no loss* cues onto context-level signal averaged over only trials that were not in the *no loss* condition. This allowed us to examine the effect of the central-parietal low-*γ* signal on behavior, dissociated from cue information. We found that this neural trace most strongly moderated RT to *no loss* cue trials in the prevention context. Furthermore, individual differences in this neural response to the loss features of the context significantly predicted behaviour to prevent loss, particularly in the *prevention* context. This provides nuance to our results by demonstrating varied patterns of behavior across individuals, and opens up interesting avenues for future research. For instance, continuing work could examine whether these effects across people hold for non-healthy populations, such as individuals that have been diagnosed with a psychiatric or neurological illness, or whether they become more pronounced.

The contrasts we specified to delineate contextual features in our neural analyses confer an additional ability to examine whether individuals demonstrate biases at the higher-order context level. Our results build on the plethora of work on loss aversion in decision making which has largely described biases at the choice or item level (e.g., Ito et al., 1998; Tversky and Kahneman, 1991; 1992). By showing that contextual loss features are prioritized, we extend the existing work on negativity biases to high-order contextual processes. We further reinforced the relevance of this bias by showing that the demarcated neural signatures reflecting emphasis on contextual loss features exert top-down influences on choice processes. Our computational modelling provides a mechanistic explanation: given that contextual features are integrated with cue information moment-by-moment in the accumulation processes, higher-level contextual biases will drive faster accumulation for the loss prevention choice. Our model further incorporates principles like divisive normalisation (Louie et al., 2013; 2015), which allow values to be compressed in a manner that captures the local context of the choice set. Interestingly, modelling unique starting positions for the promotion and prevention contexts was not necessary to capture the context-dependent behavioral effects in the empirical data.

The contextual trace across slow Θ- and Δ-frequency bands converge with previous work that suggest a maintenance mechanism for contextual information (Cashdollar et al., 2009; Hsieh and Ranganath, 2014). Indeed, processes that sustain background information, such as the outcome magnitudes of all the choice alternatives given by the current context, are necessary to explain the context-dependent behavioral error biases to control cues described above. The central-parietal low-*γ* signature similarly converges with previous work, albeit in relation to the decision processes such as evidence accumulation rather than memory maintenance *per se* (Polanía et al., 2014). That *γ*-frequency oscillations have been linked context-cue binding (Jensen et al., 2007; Nyhus and Curran, 2009) is of particular note given our computational findings where moment-by-moment integration of context and cue information within a trial can sufficiently capture the accuracy and RT context effects in the observed behavioral data. We provide evidence explicitly linking these processes to central-parietal low-*γ* neural signatures: this neural signature significantly predicts behavior both across and within participants. This finding integrates and expands beyond previous work by demonstrating that central-parietal low-*γ* neural traces are involved in context-cue binding through evidence accumulation processes.

The present research provides important insights by disentangling the effect of cues from the context in which they exist to observe how each influence decision making processes, and provide a neural and computational description of underlying mechanism. Despite these significant contributions, there remain some limitations that should be addressed by future research. First, we employed a constrained experimental paradigm to simplify and examine these target processes. Investigating contextual influences on choice in complex environments, with potentially larger numbers of relevant cues, will be necessary to further extrapolate these processes to real-word scenarios. For instance, an initial question that arises is whether the same processes hold in scenarios when multiple cues of varying relevance are presented to individuals at once. Exciting work multidimensional decision problems (Niv et al., 2015; Wunderlich et al., 2011) provide some direction. Finally, future work examining different forms of contextual information (e.g. episodic or spatial), and whether similar processes drive integration with immediately encountered cue information, will be important for understanding the generalizability of these underlying neural and computational mechanisms.

## Acknowledgements

The authors thank Cendri Hutcherson and Katherine Duncan for helpful discussions. This work was supported by a Social Sciences and Humanities Research Council of Canada (SSHRC-496650) and a Canadian Institute of Heath Research (CIHR-111257) grant to W.A.C and by a SSHRC CGS-D doctoral award to V.M.

## Author Contributions

Conceptualization, V.M. and W.A.C.; Methodology, V.M. and W.A.C.; Software, V.M.; Formal Analysis; V.M.; Investigation, V.M. and C.K.; Writing – Original Draft, V.M. and C.K.; Writing – Review & Editing, V.M., C.K., and W.A.C.; Supervision, V.M. and W.A.C.; Funding Acquisition, W.A.C.

## References

Aiken, L.S., West, S.G., and Reno, R.R. (1991). Multiple Regression (New York: SAGE).

Badre, D., and D’Esposito, M. (2009). Is the rostro-caudal axis of the frontal lobe hierarchical? Nature. Rev. Neurosci. 10,659–669.

Behrens, T.E.J., Woolrich, M.W., Walton, M.E., and Rushworth, M.F.S. (2007). Learning the value of information in an uncertain world. Nature. Neurosci. 10,1214–1221.

Bogacz, R., Brown, E., Moehlis, J., Holmes, P., and Cohen, J.D. (2006a). The physics of optimal decision making: A formal analysis of models of performance in two-alternative forced-choice tasks. Psychol. Rev. 113, 700–765.

Bornstein, A.M., Aly, M., Feng, S., Turk-Browne, N.B., Norman, K.A., and Cohen, J.D. (2017). Perceptual decisions result from the continuous accumulation of memory and sensory evidence. bioRxiv 1–27.

Botvinick, M.M. (2012). Hierarchical reinforcement learning and decision making. Curr. Opin. Neurobiol. 22, 956–962.

Brown, S.D., and Heathcote, A. (2008). The simplest complete model of choice response time: Linear ballistic accumulation. Cogn. Psychol. 57, 153–178.

Burgess, N., Maguire, E.A., and O’Keefe, J. (2002). The human hippocampus and spatial and episodic memory. Neuron 35, 625–641.

Cashdollar, N., Malecki, U., Rugg-Gunn, F.J., Duncan, J.S., Lavie, N., and Düzel, E. (2009). Hippocampus-dependent and -independent theta-networks of active maintenance. Proc. Natl. Acad. Sci. 106, 20493–20498.

Cavanagh, J.F., and Frank, M.J. (2014). Frontal theta as a mechanism for cognitive control. Trends Cogn. Sci. 18, 414–421.

Churchland, A.K., Kiani, R., and Shadlen, M.N. (2008). Decision-making with multiple alternatives. Nat. Neurosci. 11, 693–702.

Cohen, J.D., Botvinick, M., and Carter, C.S. (2000). Anterior cingulate and prefrontal cortex: who’s in control? Nat. Neurosci. 3, 421–423.

Cohen, M.X. (2014). Analyzing Neural Time Series Data: Theory and Practice (Cambridge: MIT Press).

Collins, A., and Frank, M.J. (2013). Cognitive control over learning: creating, clustering, and generalizing task-set structure. Psychol. Rev. 120, 190–229.

Daw, N.D., Gershman, S.J., Ben Seymour, Dayan, P., and Dolan, R.J. (2011). Model-Based Influences on Humans’ Choices and Striatal Prediction Errors. Neuron 69, 1204–1215.

Delorme, A., and Makeig, S. (2004). EEGLAB: an open source toolbox for analysis of single-trial EEG dynamics including independent component analysis. J. Neurosci. Methods 134, 9–21.

Diana, R.A., Yonelinas, A.P., and Ranganath, C. (2007). Imaging recollection and familiarity in the medial temporal lobe: a three-component model. Trends Cogn. Sci. 11, 379–386.

Ditterich, J., Mazurek, M.E., and Shadlen, M.N. (2003). Microstimulation of visual cortex affects the speed of perceptual decisions. Nat. Neurosci. 6, 891–898.

Dolan, R.J., and Dayan, P. (2013). Goals and Habits in the Brain. Neuron 80, 312–325.

Gershman, S.J., Blei, D.M., and Niv, Y. (2010). Context, learning, and extinction. Psychol. Rev. 117, 197–209.

Gold, J.I., and Shadlen, M.N. (2001). Neural computations that underlie decisions about sensory stimuli. Trends Cogn. Sci. 5, 10–16.

Hare, T.A., Schultz, W., Camerer, C.F., O’Doherty, J.P., and Rangel, A. (2011). Transformation of stimulus value signals into motor commands during simple choice. Proc. Natl. Acad. Sci. 108, 18120–18125.

Higgins, E.T. (1997). Beyond pleasure and pain. Am. Psychol. 52, 1280–1300.

Hsieh, L.-T., and Ranganath, C. (2014). Frontal midline theta oscillations during working memory maintenance and episodic encoding and retrieval. NeuroImage 85, 721–729.

Hunt, L.T., Kolling, N., Soltani, A., Woolrich, M.W., Rushworth, M.F.S., and Behrens, T.E.J. (2012). Mechanisms underlying cortical activity during value-guided choice. Nat. Neurosci. 15, 470–476.

Hutcherson, C.A., Bushong, B., and Rangel, A. (2015). A Neurocomputational Model of Altruistic Choice and Its Implications. Neuron 87, 451–462.

Ito, T.A., Larsen, J.T., Smith, N.K., and Cacioppo, J.T. (1998). Negative information weighs more heavily on the brain: the negativity bias in evaluative categorizations. J. Pers. Soc. Psychol. 75, 887–900.

Jensen, O., Kaiser, J., and Lachaux, J.-P. (2007). Human gamma-frequency oscillations associated with attention and memory. Trends Neurosci. 30, 317–324.

Krajbich, I., and Rangel, A. (2011). Multialternative drift-diffusion model predicts the relationship between visual fixations and choice in value-based decisions. Proc. Natl. Acad. Sci. 108, 13852–13857.

Louie, K., Grattan, L.E., and Glimcher, P.W. (2011). Reward Value-Based Gain Control: Divisive Normalization in Parietal Cortex. J. Neurosci. 31, 10627–10639.

Louie, K., Glimcher, P.W., and Webb, R. (2015). Adaptive neural coding: from biological to behavioral decision-making. Curr. Opin. Behav. Sci. 5, 91–99.

Louie, K., Khaw, M.W., and Glimcher, P.W. (2013). Normalization is a general neural mechanism for context-dependent decision making. Proc. Natl. Acad. Sci. 110, 6139–6144.

MacDonald, A.W., Cohen, J.D., Stenger, V.A., and Carter, C.S. (2000). Dissociating the Role of the Dorsolateral Prefrontal and Anterior Cingulate Cortex in Cognitive Control. Science 288, 1835–1838.

Mensen, A., and Khatami, R. (2013). Advanced EEG analysis using threshold-free cluster-enhancement and non-parametric statistics. NeuroImage 67, 111–118.

Mognon, A., Jovicich, J., Bruzzone, L., and Buiatti, M. (2011). ADJUST: An automatic EEG artifact detector based on the joint use of spatial and temporal features. Psychophysiology 48, 229–240.

Mormann, M.M., Malmaud, J., Huth, A., Koch, C., and Rangel, A. (2010). The Drift Diffusion Model Can Account for the Accuracy and Reaction Time of Value-Based Choices Under High and Low Time Pressure. Judgm. Decis. Mak. 5, 437–449.

Mowrer, S.M., Jahn, A.A., Abduljalil, A., and Cunningham, W.A. (2011). The Value of Success: Acquiring Gains, Avoiding Losses, and Simply Being Successful. PLoS ONE 6, e25307–e25308.

Mulder, M.J., Wagenmakers, E.J., Ratcliff, R., Boekel, W., and Forstmann, B.U. (2012). Bias in the Brain: A Diffusion Model Analysis of Prior Probability and Potential Payoff. J. Neurosci. 32, 2335–2343.

Nieuwenhuis, S., Holroyd, C.B., Mol, N., and Coles, M.G.H. (2004). Reinforcement-related brain potentials from medial frontal cortex: origins and functional significance. Neurosci. Biobehav. Rev. 28, 441–448.

Niv, Y., Daniel, R., Geana, A., Gershman, S.J., Leong, Y.C., Radulescu, A., and Wilson, R.C. (2015). Reinforcement Learning in Multidimensional Environments Relies on Attention Mechanisms. J. Neurosci. 35, 8145–8157.

Nyhus, E., and Curran, T. (2009). Semantic and perceptual effects on recognition memory: Evidence from ERP. Brain Res. 1283, 102–114.

Oostenveld, R., Fries, P., Maris, E., and Schoffelen, J.-M. (2011). FieldTrip: Open Source Software for Advanced Analysis of MEG, EEG, and Invasive Electrophysiological Data. Comput Intell. Neurosci. 2011, 1–9.

Polanía, R., Krajbich, I., Grueschow, M., and Ruff, C.C. (2014). Neural oscillations and synchronization differentially support evidence accumulation in perceptual and value-based decision making. Neuron. 82, 709–720.

Ratcliff, R. (1978). A Theory of Memory Retrieval. Psychol. Rev. 85, 59–108.

Ribas-Fernandes, J.J., Solway, A., Diuk, C., McGuire, J.T., Barto, A.G., Niv, Y. and Botvinick, M.M. (2011). A neural signature of hierarchical reinforcement learning. Neuron, 71, 370–379.

Roe, R.M., Busemeyer, J.R., and Townsend, J.T. (2001). Multialternative decision field theory: a dynamic connectionist model of decision making. Psychol. Rev. 108, 370–392.

Rushworth, M.F.S., Noonan, M.P., Boorman, E.D., Walton, M.E., and Behrens, T.E. (2011). Frontal Cortex and Reward-Guided Learning and Decision-Making. Neuron 70, 1054–1069.

Schuck, N.W., Cai, M.B., Wilson, R.C., and Niv, Y. (2016). Human Orbitofrontal Cortex Represents a Cognitive Map of State Space. Neuron 91, 1402–1412.

Shadlen, M.N., and Newsome, W.T. (2001). Neural basis of a perceptual decision in the parietal cortex (area LIP) of the rhesus monkey. J. Neurophysiol. 86, 1916–1936.

Smith, S.M., and Nichols, T.E. (2009). Threshold-free cluster enhancement: addressing problems of smoothing, threshold dependence and localisation in cluster inference. NeuroImage 44, 83–98.

Solway, A., and Botvinick, M.M. (2012). Goal-directed decision making as probabilistic inference: A computational framework and potential neural correlates. Psychol. Rev. 119, 120–154.

Summerfield, C., and Mangels, J.A. (2005). Coherent theta-band EEG activity predicts item-context binding during encoding. NeuroImage 24, 692–703.

Tavares, G., Perona, P., and Rangel, A. (2017). The Attentional Drift Diffusion Model of Simple Perceptual Decision-Making. Front. Neurosci. 11, 396–16.

Tolman, E.C. (1948). Cognitive maps in rats and men. Psychol. Rev. 55, 189–208.

Tse, D., Langston, R.F., Kakeyama, M., Bethus, I., Spooner, P.A., Wood, E.R., Witter, M.P., and Morris, R.G.M. (2007). Schemas and memory consolidation. Science 316, 76–82.

Tsetsos, K., Usher, M., and McClelland, J.L. (2011). Testing multi-alternative decision models with non-stationary evidence. Front. Neurosci. 5, 63.

Tversky, A., and Kahneman, D. (1991). Loss Aversion in Riskless Choice: A Reference-Dependent Model. Q. J. Econ. 106, 1039–1061.

Tversky, A., and Kahneman, D. (1992). Advances in prospect theory: cumulative representation of uncertainty. J. Risk Uncertain. 5, 297–323.

Tversky, A., and Simonson, I. (1993). Context-dependent Preferences. Management Science 39, 1179–1189.

Usher, M., and McClelland, J.L. (2001). The time course of perceptual choice: the leaky, competing accumulator model. Psychol. Rev. 108, 550–592.

Wald, A. (1947). Foundations of a General Theory of Sequential Decision Functions. Econometrica 15, 279–313.

Wilson, R.C., Takahashi, Y.K., Schoenbaum, G., and Niv, Y. (2014). Orbitofrontal Cortex as a Cognitive Map of Task Space. Neuron 81, 267–279.

Wunderlich, K., Beierholm, U.R., Bossaerts, P., and O’Doherty, J.P. (2011). The human prefrontal cortex mediates integration of potential causes behind observed outcomes. J. Neurophysiol. 106, 1558–1569.

